# Arabidopsis exocyst complex subunit EXO70E2 in defence against *Pseudomonas syringae* in conjunction with autophagy

**DOI:** 10.64898/2026.06.30.735562

**Authors:** Ali Burak Yıldız, Andrea Potocká, George Alexandru Caldarescu, Adam Batík, Peter Sabol, Viktor Žárský

## Abstract

Exocyst was initially uncovered in yeast genetic *sec*-screen as a tethering complex for exocytotic vesicles and this function was later found to be evolutionarily conserved in other eukaryotes including plants. Later however, a surprising engagement of the exocyst complex in autophagy was observed in animals, plants and recently also in yeast. Using the genetic approach we observed EXO70E2 exocyst complex subunit engagement in the defence response to *Pseudomonas syringae* attack linked to the autophagy pathway. CRISPR/CAS LOF mutant of EXO70E2 is more sensitive to *Pseudomonas* infection (both virulent as well as T3SS mutant) and autophagy flux monitored by NBR1 antibody is compromised in comparison to WT. We conclude that the plant exocyst complex linked to the EXO70E2 subunit participates in defence against *Pseudomonas* bacteria in conjunction with the autophagy pathway.

**Highlight:** Arabidopsis exocyst subunit EXO70E2 affects selective autophagic flux monitored by NBR1 and is participating in defense against *Pseudomonas syringae* infection.

## Introduction

The exocyst is a conserved multi-subunit protein complex found in all eukaryotes. It comprises eight subunits: SEC3, SEC5, SEC6, SEC8, SEC10, SEC15, EXO70 and EXO84, which form a hetero-oligomeric complex (TerBush *et al*., 1996). Exocyst functions as a CATCHR-type (complexes associated with tethering containing helical rods) vesicle-tethering complex and as an effector of RAB and RHO GTPases plays a crucial role in the tethering and docking of post-Golgi secretory vesicles to the plasma membrane (PM) and regulation of soluble *N-ethylmaleinide-*sensitive-factor (SNARE) complex formation, resulting in the vesicle fusion at the PM. Also, the plant exocyst complex is divided into two modules: the first, composed of SEC3, SEC5, SEC6, and SEC8; and the second, composed of SEC10, SEC15, EXO70, and EXO84 (Synek *et al*., 2021). In land plants, the EXO70 subunit family exhibits a unique pattern of expansion and divergence, giving rise to three subfamilies (EXO70.1, EXO70.2, EXO70.3) with rapidly evolving paralogues within EXO70.2 subfamily partly due to the “Red Queen” like competition between the hosts and parasites (Synek *et al*., 2006; Cvrčková *et al*., 2012; Zárský *et al*., 2013; Žárský *et al*., 2020; Holden *et al*., 2022). The functional significance of specific EXO70 isoforms however relate not only to cellular defence and innate immunity. EXO70B1 and EXO70B2 play pivotal roles in PAMP-triggered immunity (the membrane localisation of PRRs, callose deposition, cell wall modifications; (Pecenková *et al*., 2011; Stegmann *et al*., 2012; Wang *et al*., 2020; Ortmannová *et al*., 2022; Drs *et al*., 2025). In grasses, the rapidly evolving specific FX clade of EXO70s diversifies in response to the pathogens pressure (Holden *et al*., 2022). Also core exocyst subunits are required for plant immunity (Du *et al*., 2015, 2018; Michalopoulou *et al*., 2022). But e.g. EXO70H4 linked exocyst is involved in localized deposition of callose in the upper domain of Arabidopsis single cell trichomes (Kulich *et al*., 2015, 2018).

On top of that, it has been found that the EXO70B1-containing version of exocyst complex is important for import of secondary metabolites (esp. anthocyanins) into the vacuole via autophagy-related membrane containers (Kulich *et al*., 2013). This corroborated previous discovery of exocyst engagement in autophagosome initiation in metazoa (Bodemann *et al*., 2011) and opened new unexpected perspectives on exocyst cellular functions not only in the conventional exocytosis, but also in processes related to the autophagy (Pecenková *et al*., 2017b). The closest homolog of EXO70B1, EXO70B2, is transported to the vacuole upon phosphorylation, possibly for its subsequent degradation (Teh *et al*., 2018) and PM localized EXO70B2 is re-directed into the autophagy pathway upon phosphorylation (Brillada *et al*., 2021). Similarly, EXO70D isoforms were shown to play a major part in specific/targeted autophagic degradation of Type-A ARR proteins to regulate cytokinin sensitivity as novel specific autophagy receptors (Acheampong *et al*., 2020). Importantly, both EXO70B1 and B2 co-localize with ATG8 autophagosomal maker, possibly related to the presence of ATG8-interacting motifs (AIMs; Cvrčková and Zárský, 2013; Tzfadia and Galili, 2013; Teh *et al*., 2018). Also, the SEC5 exocyst subunit has been implicated in autophagy regulation through interaction with ROP8, a member of the plant-specific branch of Rho GTPases (Lin *et al*., 2021).

Macroautophagy (hereafter autophagy) is a conserved intracellular degradation and recycling pathway essential for cellular homeostasis, nutrient recycling, and stress adaptation across eukaryotes (Marshall and Vierstra, 2018). Initiation of autophagy occurs with the *de novo* formation of a double-membrane vesicle called the phagophore, which expands to engulf cytoplasmic components, protein aggregates, or damaged organelles. This vesicle closes to form a mature autophagosome, which subsequently targets and fuses with the vacuole or the lysosome (Marshall and Vierstra, 2018). But initially, autophagy was considered an unrestricted degradation machinery; by employing various receptors, it can recognise specific cargos to tether into forming autophagosome to be degraded (Farré and Subramani, 2016; Dikic, 2017; Marshall and Vierstra, 2018). In plants, this selectivity relies on the cargo receptor Neighbour of BRCA1 1 (NBR1), a functional hybrid that shares structural domains with the mammalian receptors p62/SQSTM1 and NBR1 (Svenning *et al*., 2011). While it can label polyubiquitinated protein aggregates for degradation thanks to its Ubiquitin-Associated (UBA) domain, it can also directly capture ATG8 proteins via its ATG8-Interacting Motif (AIM) domain (Svenning *et al*., 2011; Zhou *et al*., 2014). In addition to ubiquitin-dependent selection, NBR1 can isolate non-ubiquitinated cargo via liquid-liquid phase separation (LLPS), forming condensed liquid droplets that serve as specialized templates for autophagosome nucleation (Zaffagnini *et al*., 2018; Yan *et al*., 2024). During pathogen invasion or abiotic stress, NBR1 is involved in selecting specific structural or regulatory proteins and directing them to the vacuole for degradation (Hafrén *et al*., 2018; Yan *et al*., 2024). Since the 2010 report (Wang *et al*., 2010) EXO70E2 paralog has been studied and discussed, due to effects of its ectopic overexpression, as marker of specific double membrane containers, not related to the autophagy, and secreted to the apoplast in an unconventional manner (“EXPO”; also Ding *et al*., 2014; Lin *et al*., 2015). Further studies from this lab have shown that due to direct interaction between EXO70E2 and autophagy receptor NBR1 the double membrane EXO70E2 positive compartment may be encased by the double membrane autophagosome (Ji *et al*., 2020). Studies on unconventionally secreted apoplastic vesicles established EXO70E2 positive population as a well-defined type of extracellular vesicles (Koch *et al*., 2026). Yan et al. (2024) based on a deep mechanistic analysis of EXO70E2 involvement with the autophagy unequivocally demonstrated that EXO70E2 is a non-ubiquitinated substrate of a selective autophagy mediated by liquid-liquid phase separation feature of plant NBR1, so that EXO70E2 positive bodies in plant cells include NBR1 phase separated droplets with aggregated EXO70E2 (or other substrates) which are encased by the phagophore in a next phase (Yan *et al*., 2024). There is also hint that EXO70E2 function is linked to the ESCRT complex via ISTL1; ESCRT complex has a possible role in autophagosome closure (Camacho, 2019; Zhou *et al*., 2019); both supporting possibility of EXO70E2 itself functioning in autophagy.

Thus, it becomes necessary to understand how autophagy - and especially NBR1-mediated autophagy - is affected by the EXO70E2. fosWhile studies on NBR1-EXO70E2 relation are based mostly on cell biological observations using *nbr1* LOF mutants, there are only few reports on attempts to study EXO70E2 functional implications using *exo70E2* LOF mutants (first T-DNA and recently also CRISPR; (Neves *et al*., 2021, 2022; Copeland *et al*., 2025; Koch *et al*., 2026). Here we report a functional analysis of an *exo70E2* CRISPR LOF allele created by us. In accordance with expression data (ePlant, University of Toronto, Waese *et al*., 2017; Pečenková *et al*., 2020), the EXO70E2 was found to be involved in the biotic interaction of Arabidopsis with *Pseudomonas syringae* pathogen - we observed induction of EXO70E2 accumulation after the pathogene attack and an enhanced sensitivity to the pathogen and partially compromised selective NBR1-monitored autophagy flux of the *exo70E2* mutant.

## Material and methods

### Plant Growth and Conditions

The *in vitro* cultivation of Arabidopsis plants was carried out at 22°C, the light was provided by LED light tubes (T8-HBN120-DW, 4000-4500 K), in a long-day setting with a photoperiod of 16/8-h ight/darkness. Light intensity was 90-110 mol/s/m^2^. Except the plant material for callose staining, all the plant material was cultivated in a long-day setting with a photoperiod of 16/8-h light/darkness was carried out at 22°C/19°C (day/night). Light was provided by LED strips producing around 110 mol/s/m^2^. For the callose staining, Arabidopsis plants were grown in the cultivation room under the short-day settings with a photoperiod of 10/14-h light/darkness.

### Generation of plant expression constructs and transgenic lines

To generate *exo70E2* CRISPR mutants, 2 guide RNAs (gRNAs) carrying a CRISPR cassette were constructed according to a previously published multiplex gene-editing method (Xing *et al*., 2014). CRISPR-P web tool (Lei *et al*., 2014) was used to design gRNAs. gRNAs were then amplified together with the target fragments from pCBC-DT1DT2 (https://www.addgene.org/50590/). The cassette was then cloned into pHSE401 (https://www.addgene.org/62201/) using Golden Gate cloning (Engler *et al*., 2008). The final vector with the double gRNA cassette was transferred into *Agrobacterium tumefaciens* (strain GV3101) (Koncz and Schell, 1986) and then used in the floral dip method to transform *A. thaliana* Col-0 plants (Clough and Bent, 1998). After the transformation, plants were selected on hygromycin plates, and the point mutations were validated via Sanger DNA sequencing. For the complementation of the *exo70E2* mutant, the pEXO70E2::GFP::EXO70E2 construct was generated via Gateway cloning and cloned into pB7m34GW (https://vectorvault.vib.be/collection/pb7m34gw0) and final vector was transferred into the *Agrobacterium tumefaciens* (GV3101). Then *Agrobacterium* transformed into transgenic *exo70E2* CRISPR mutant plants via the floral dip method. Transgenic plants were selected on Basta (Glufosinate ammonium, Sigma – Aldrich) selective plates.

### Bacterial Infection and Resistance Assay

The flood-inoculation protocol for Arabidopsis seedlings was adapted from Ishiga *et al*. (Ishiga *et al*., 2011). Seeds were sterilised in 1.5 ml Eppendorf tubes, starting with 3 mins of washing with 70% ethanol, followed by two washes with 10% bleach for 3 mins each. Washed three times with sterile distilled water and kept for two days at 4°C to break the dormancy. Plated on horizontal round ½ MS plates, 0.8% plant agar, deep Petri plates; 12 seedlings per plate. Then, seedlings were grown for 11 days under a 12h light/12h dark photoperiod. Pre-infection fresh *Pseudomonas syringae pv. tomato DC3000* and *hrph-* overnight plate culture was prepared on Luria-Bertani low salt (LBLS) medium containing appropriate antibiotics. For flood inoculation, a sterile 0.0025% Silwet L-77 solution was prepared to resuspend the bacteria. For the *DC3000* strain, OD_600_ = 0.01, and the hrph- strain, OD_600_ = 0.05, were used. Approximately 40-50 ml of solution was poured into each plate and incubated for 2 minutes at room temperature. After the incubation, the flooding suspension was removed from the plates, and the plates were moved back to the growth room. On the 1st day post inoculation (dpi), the internal bacterial populations in each replicate were evaluated from 4 pooled independent seedlings grown in 3 Petri dishes. Each sample was weighed pre-extraction and washed for 20 seconds in 70% ethanol, followed by 10 seconds in sterile distilled water. Samples were homogenized using the Omni Bead Ruptor Elite (Omni International) for 30 seconds at 4 m/s. Serial dilutions were made up to 10^-6^, and plated onto LBLS with corresponding antibiotics, and grown at 28 degrees for 24 hours. Measured weights were used to normalize bacterial growth. The same seedlings were also used for immunoblotting analysis, and for the control, 0.0025% Silwet L-77 water solution without any bacteria was used.

The root tip droplet assay for Arabidopsis seedlings was adapted from Pečenková et al. ^(^Pecenkov^á^ *et al*., 2017*a*). We only adjusted the OD_600_ for *Pst DC3000* to 0.1 and infected as per the published protocol. 4 dpi, fresh weights were measured with hypocotyls.

### Callose Staining and deposition

According to the growth spiral, the 7th, 8th, and 9th leaves of 5-week-old Arabidopsis plants were infiltrated with *Pst hrph-* at an OD_600_ of 0.1. 24 hpi, infected leaves were harvested and washed for 3 h in acetic acid:ethanol (1:3), followed by three washes with water and incubation in aniline blue solution (150 mM KH_2_PO_4_ and 0.01% [w/v] aniline blue, pH 9.5) (Kulich *et al*., 2015). After the staining, the whole leaves were scanned, and 4 to 6 ROI per leaf were generated. The number of callose per ROI was measured via Dot Scanner (Allen *et al*., 2024).

### E-64d treatment of Arabidopsis seedlings and area coverage analysis

E-64d/FM4-64 treatment was adopted from Goto-Yamada et al.(Goto-Yamada *et al*., 2019), with some changes. 5-day-old Arabidopsis seedlings grown on growth media were transferred to liquid 1/2 media without sucrose, containing 4 µM FM4-64 and 5 µM E-64d and covered with foil to keep them under dark conditions, and root cells were observed after 24h under a spinning disc microscope. To cover the entire root, multiple Z-stacks were made from the root tip to the mature zone of each root. A total of 100 cells were picked for each genotype, and 11 layers of z-stack were created for each cell. Then, with ImageJ, the maximum intensity projection of these cells was created and measured by a self-written macro to analyse the total area covered by E-64d bodies per cell.

### Immunoblot analysis

Twelve-day-old Arabidopsis seedlings (appx 30-40 mg) were ground in liquid nitrogen. After the grounding, 250 μL of Sec6/8 buffer (20 mM HEPES, pH 6.8, 150 mM NaCl, 1 mM EDTA, 1 mM DTT, and 0.5% Tween 20) was added and supplemented with 1× protease inhibitor cocktail (Sigma-Aldrich) (Hála *et al*., 2008). The total lysate was then centrifuged at 1000g for 10 min. Extracts were boiled in the SDS loading buffer. Samples were loaded onto 12-well 4-15% criterion TGX Stain-Free gels (Bio-Rad, Hercules, CA) and followed by 200V 30 min electrophoresis. Next, Stain-Free gel activation was performed for 45 seconds with the Bio-Rad Chemidoc imaging system. After the activation, the gel was transferred onto a mini size membrane by using a Trans-Blot Turbo transfer system for 7 min (Bio-Rad, Hercules, CA). Membranes were blocked with 5% skimmed milk in PBS, and incubated with primary antibody anti-NBR1 (Agrisera) using 1:4000 dilutions in PBS containing 0.1% Tween 20 overnight, followed by Rabbit horseradish peroxidase-conjugated secondary antibodies diluted 1:10,000 in PBS containing 0.1% Tween 20 for 55 minutes. The ECL prime kit was used to develop and visualise with Bio-Rad ChemiDoc Imaging Systems (Bio-Rad, Hercules, CA).

### Microscopy

To validate complementation of *exo70E2* CRISPR mutants and measure the formation of E-64d bodies, imaging was performed using a vertical stage Zeiss Axio Observer 7 coupled to a Yokogawa CSU-W1-T2 spinning disk unit with 50 µm pinholes and equipped with a VS-HOM1000 excitation light homogenizer (Visitron Systems) (von Wangenheim *et al*., 2017; Serre *et al*., 2021). We used Zeiss Plan-Apochromat ×20/0.8 M27 (FWD=0.55mm) and alpha Plan-Apochromat 100x/1.46 Oil DIC M27 (FWD=0.10mm), (UV)VIS-IR and acquisition of images was done by VisiView software (Visitron Systems, v.4.4.0.14). The FM4-64-stained samples were excited by 561 nm laser, emission to 582 to 636 nm and for GFP 488 nm laser used with emission 500 to 550 nm (Kubalová *et al*., 2025). The signal was detected using a PRIME-95B Back-Illuminated sCMOS camera (1,200 × 1,200 pixels; Photometrics).

For the acquisition of callose deposition experiment, Zeiss Axiolmager with ApoTome2 module was used. Images of whole leaves were taken by objective EC Plan-Neofluar 5x/0.16 M27 and samples were excited by 353 nm laser (DAPI).

### Statistical Analysis and Data Imputation

R v4.5.1 (R foundation, https://www.r-project.org/) was used for plotting, statistical analyses, and filtering of some data during this study. Plotting was carried out using the ggplot2 library, whereas data processing was facilitated by the *dplyr* library, included in the Tidyverse package (Wickham *et al*., 2019).

## Results

### *exo70E2* mutant is more sensitive to both WT and TTSS mutant *Pseudomonas syringae* infection and this correlates with the defect in callose deposition

To address function of EXO70E2 Arabidopsis paralog genetically we created a new CRISPR/CAS LOF allele of this single exon locus inducing a frame-shift proximal to the 5’- end of the coding sequence (Fig. 1 A; *exo70E2*). Sequence alignment comparing the mutant line against the wild-type (WT) reference sequence revealed successful editing at the target locus, with two base pair deletions (CDS nucleotides 99 and 100; see Fig. 1A) resulting in a premature stop codon (PSC) at the marked positions. This mutant line was stably transformed with a pEXO70E2::GFP::EXO70E2 expression cassette, driving the expression of a green fluorescent protein (GFP)-tagged variant under its native promoter. Transgenic Arabidopsis plant roots revealed a distinct and strong fluorescent signal localised within the root cap cells, similarly to GUS staining in the root tip observed by Wu and Huang (Wu and Huang, 2026). Despite the expression maxima of this gene in root cap (Fig. 1B; supported by ePlant Toronto Arabidopsis gene expression database) we did not observe any defects in root cap development in *exo70E2* mutant and also recapitulation of some published stress treatments (osmotic; Neves *et al*., 2022) did not show any phenotypic deviation of this mutant as compared to WT (Fig. S1). However, ePlant EXO70E2 expression data indicate distinct expression also in leaves and induction of EXO70E2 mRNA in response to pathogen or elicitor treatments. We therefore used *Pseudomonas syringae* bacterial pathogen (both virulent pv. *tomato DC3000* and T3SS mutant *hrph-*) to test possible change of *exo70E2* mutant sensitivity to the infection. 11-day-old seedlings of wild-type (WT), the *exo70E2* CRISPR mutant, and the *exo70E2* complemented lines were subjected to flood inoculation assays. Seedlings were challenged with either the fully virulent pathogen *Pseudomonas syringae pv. tomato (Pst) DC3000* or type III secretion system (T3SS)-deficient mutant *Pst hprh^-^*. Following inoculation with virulent *Pst DC3000*, the *exo70E2* mutant line exhibited significantly increased susceptibility compared to WT plants, as evidenced by enhanced bacterial growth (Figure 1C). This phenotype was successfully rescued in complemented lines expressing the pEXO70E2::GFP::EXO70E2 construct, in which bacterial growth and susceptibility were restored to WT levels (Figure 1C). When challenged with the T3SS compromised *Pst hprh^-^* strain, *exo70E2* mutant likewise supported significantly greater bacterial growth than WT plants (*** p<0.001) indicating enhanced susceptibility to this strain as well. We therefore observed that loss of EXO70E2 increased susceptibility to both of the pathogen strains and this phenotypic deviation was successfully complemented by pEXO70E2::GFP::EXO70E2 expression cassette – indicating also biological functionality of this construct. Consistent with this phenotype, infection also resulted in a reduction of fresh weight in *exo70E2* mutant compared to the Col-0 WT plants (Fig. 2C).

**Figure 1.**
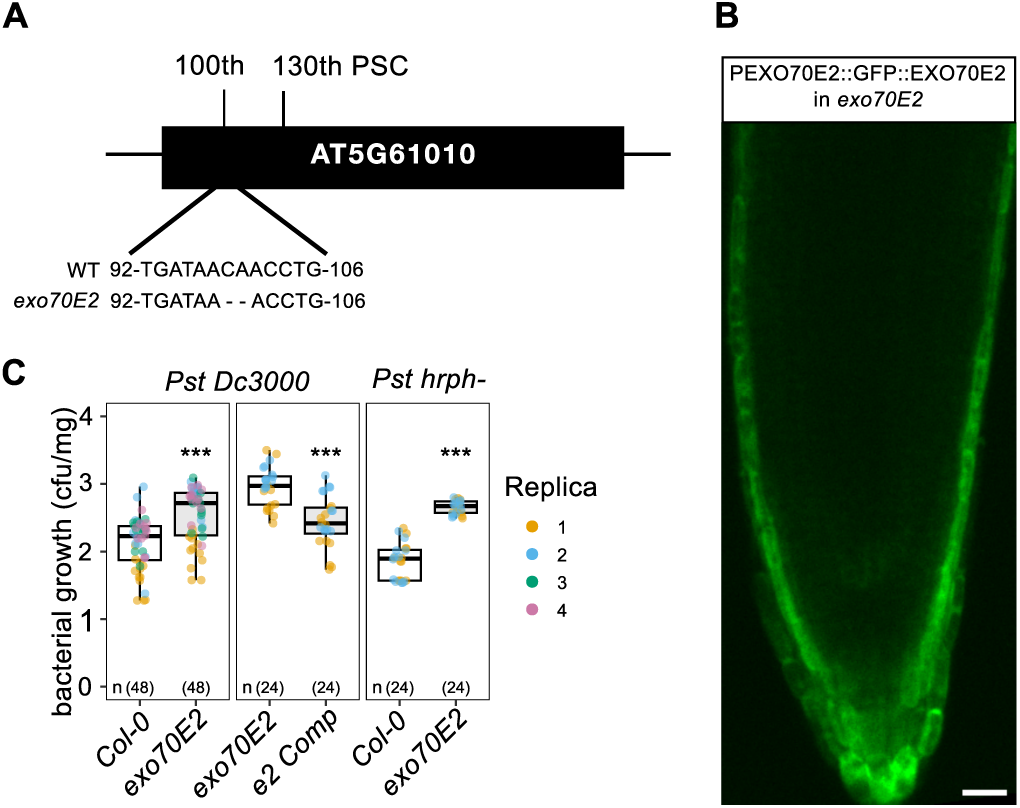
Characterization of the *exo70E2* CRISPR/Cas9 mutant and complementation of the mutant with the pEXO70E2::GFP::EXO70E2 expression cassette (hereafter e2 Comp). (A) CRISPR/Cas9-Induced mutation analysis of EXO70E2 (AT5G61010). Reference sequence revealed successful editing at the target locus, with two base pair deletions resulting in a premature stop codon (PSC) at the marked positions. (B) Subcellular Localization and Complementation of EXO70E2 with a pEXO70E2::GFP::EXO70E2 expression cassette to confirm that the disruption of EXO70E2 specifically caused the observed mutant phenotypes and root cap of 7-day-old Arabidopsis plants were used to observe localization of EXO70E2 (scale bar = 20 µm). (C) Pathogen susceptibility and complementation analysis to evaluate the role of EXO70E2 in plant immunity, 11-day-old seedlings of wild-type (WT), the *exo70E2* mutant, and the e2 Comp lines were subjected to flood inoculation assays. Seedlings were challenged with either *Pst DC3000* or *Pst hprh^-^*for 24 hours(*** p<0.001; linear mixed-effects model with genotype as a fixed effect and biological replicate as a random effect; post hoc pairwise comparisons were performed using estimated marginal means).

**Figure 2.**
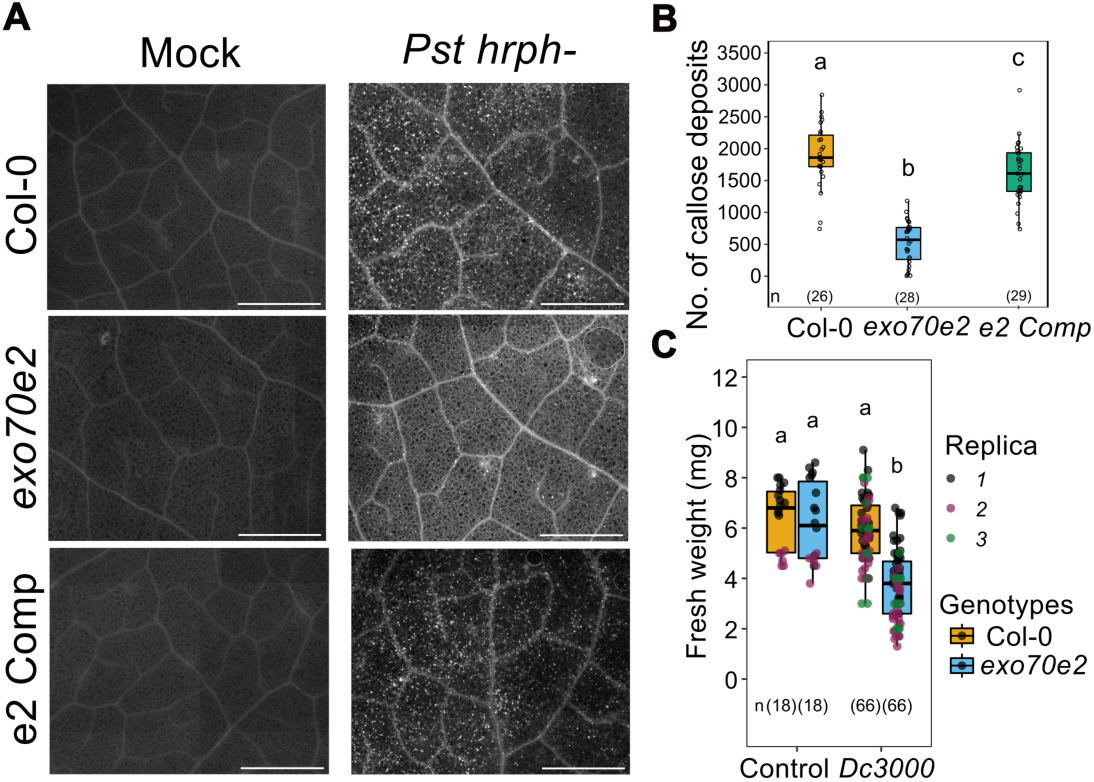
Evaluation of Callose Deposition and Susceptibility to *Pseudomonas syringae* pv. *tomato* (*Pst*). (A) Callose deposition in Arabidopsis leaves. Representative micrographs of aniline blue-stained leaves from Col-0, *exo70E2* mutant, and e2 Comp line at 24 hours post-infiltration with either a mock solution (left panels) or *Pst* hrph- (right panels). Scale bars = 1 mm. (B) Quantification of callose deposition. The number of callose foci per ROI (ROI area = 9.15 mm^2^) following *Pst* hrph- across Col-0 (yellow), *exo70E2* (blue), and complementation lines (green). Centre lines represent the median, box limits indicate the 25th and 75th percentiles, and individual data points are shown as open circles (n = X ROI per genotype). Distinct letters indicate statistically significant differences (p < 0.05, ANOVA with Tukey’s HSD test). (C) Plant growth and susceptibility assay after *Pst* DC3000 infection. Fresh weight of 10-day-old Arabidopsis seedlings measured 4 days post-inoculation (dpi) with a 10 μl droplet of *Pst* DC3000 (OD_600_ = 0.1). Data points are colour-coded by replicate, comparing Col- 0 and *exo70E2* mutants under mock (water droplets) or infection conditions. Boxplots show the distribution of fresh weights and horizontal line shows median value (n = X seedlings per group). Different letters denote statistical differences as determined by Two-way ANOVA with HSD test.

As callose deposition is a known process related to plant defence against pathogens we stained control and *Pseudomonas* infected leaves by aniline blue chromophore and observed significantly decreased callose deposition in *exo70E2* as compared to the WT (Fig. 2A). This deviation is also rescued in complemented lines (Fig. 2B). These observations clearly imply EXO70E2 exocyst subunit in processes of defence against *Pseudomonas* bacterial pathogen.

### *Pseudomonas* infection-related NBR1-dependent autophagy flux is partially compromised in *exo70E2* LOF mutant

As EXO70E2 exocyst subunit has been implicated in the NBR1-dependent autophagosome loading as a cargo (Ji *et al*., 2020; Yan *et al*., 2024) and *Pseudomonas* manipulates autophagy in infected plants (Mostowy, 2013; Üstün *et al*., 2018), we investigated whether autophagic flux in infected WT and *exo70E2* mutant plants by monitoring autophagy following vacuolar alkalinization with an inhibitor of V-ATPase Concanamycin A (ConA). Seedlings were subjected to flood inoculation with either virulent (Fig. 3A) *Pst DC3000* or (Fig. 3B) *Pst hprh^-^*, in the presence (+) or absence (–) of ConA. Both *Pseudomonas* strains infections induced autophagic flux in WT, whereas this activation was largely compromised in the *exo70E2* mutant (Fig. 3, A and B). Moreover, infection with *Pseudomonas syringae* enhanced protein levels of both the NBR1 and GFP-EXO70E2 (Fig. S1), as predicted by the transcriptomes (ePlant University of Toronto) further highlighting their role in defence.

**Figure 3.**
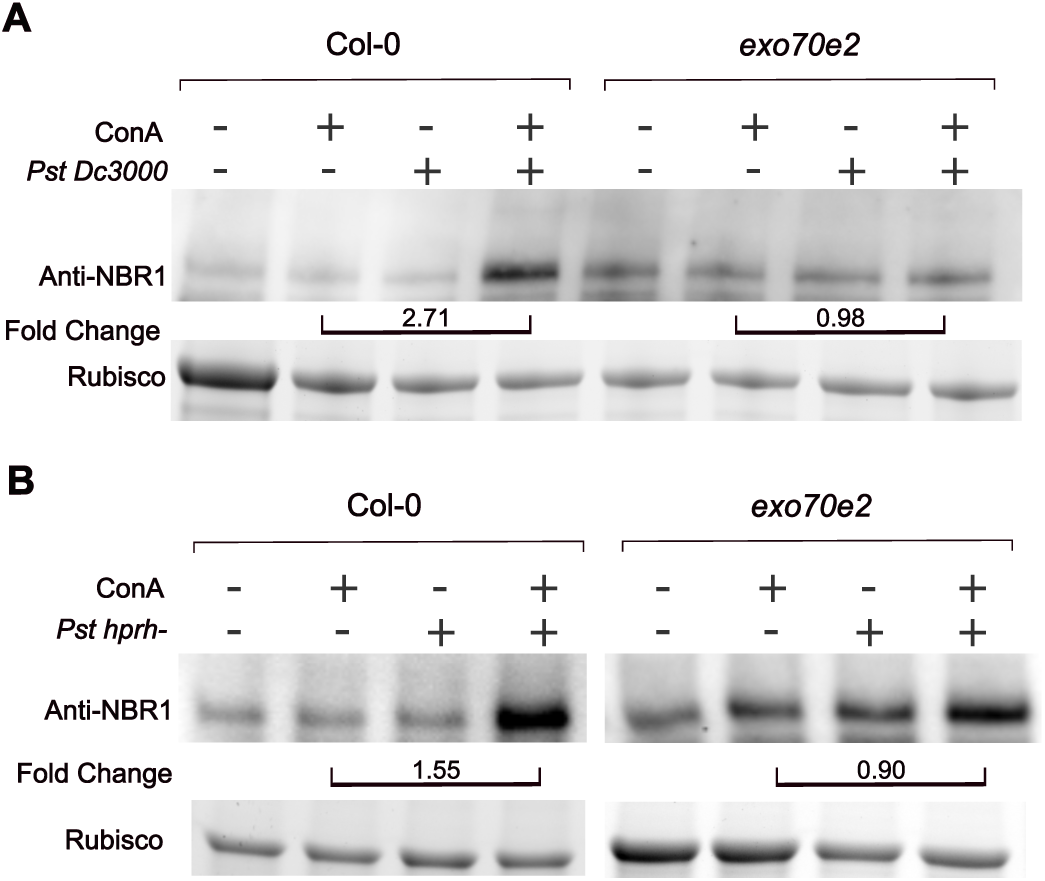
*exo70E2* mutants exhibit impaired autophagy flux. Western blot analysis of NBR1 protein accumulation and autophagic flux in 11-day-old wild-type (Col-0) and *exo70E2* mutant *Arabidopsis* seedlings. Seedlings were subjected to flooding inoculation with either virulent (A)*Pst DC3000* or (B) the type III secretion system-deficient mutant *Pst hprh–*, in the presence (+) or absence (–) of the vacuolar H⁺-ATPase inhibitor ConA at 10 hpi for additional 14h. Total protein extracts were immunoblotted with anti-NBR1 antibodies overnight. Rubisco (Bio-Rad stain-free gel activation) serves as the loading control. Fold-change values indicate the ratio of NBR1 band intensity between ConA(+) conditions for each respective genotype and treatment.

### *exo70E2* mutant shows impaired basic autophagic flux under the E-64d protease inhibitor treatment in roots

E-64d is commonly used in plant autophagy experiments as a vacuolar cysteine protease inhibitor which causes, by blocking degradation inside the vacuole, autophagic bodies to accumulate making the intensity of autophagy process easier to visualize and detect using microscopy. In agreement with above autophagy flux observations, WT Arabidopsis plant roots accumulated about 20% of cellular area autophagic bodies visualized using extended FM4-64 labelling in comparison to less than 15% in *exo70E2* mutant after the E-64d treatment (Fig. 4B and C). Comparable reduction in NBR1 signal is detected in *exo70E2* mutant by Western blot analysis (Fig. 4A). These results also indicate that EXO70E2 is positively involved in the NBR1-mediated basal autophagy visualized by E-64d-sensitive autophagosomes accumulation. Collectively, these observations of autophagy fluxes after ConA or E-64d treatments demonstrate that EXO70E2 positively modulates NBR1-associated autophagic flux.

**Figure 4.**
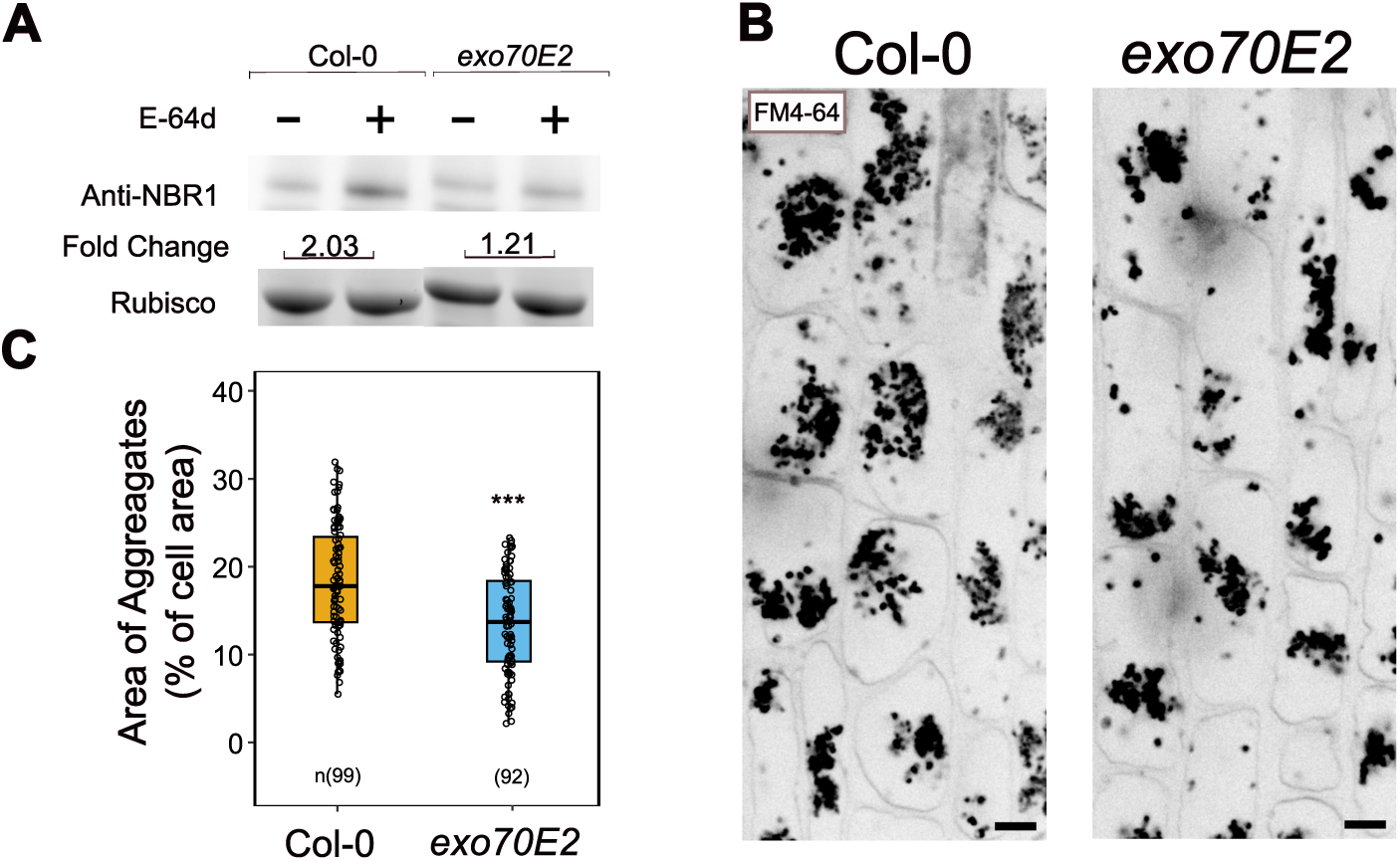
EXO70E2 deficiency impairs E-64d-sensitive autophagic flux and reduces the accumulation of vacuolar E-64d bodies. (A) Western blot analysis of NBR1 autophagic flux in 11-day-old wild-type (Col-0) and *exo70E2* mutant seedlings in the presence (+) or absence (–) of the cysteine protease inhibitor E-64d for 14h. Rubisco serves as a loading control. Fold-change values indicate the ratio of NBR1 band intensity between E-64d(+) and E-64d(–) conditions. (B) Spinning disc microscopy images of Arabidopsis root cells. Seedlings were co-treated with the lipophilic endocytic tracer FM4-64 and E-64d for 24 hours to visualise the internal accumulation of endocytic and autophagic cargo aggregates within the vacuole. (Scale bars = 5 μm). (C) Quantification of the total area of accumulated E-64d bodies normalised to total cell area (%) in Col-0 (orange) and *exo70E2* (blue) root cells following the 24-hour E-64d/FM4-64 co-treatment. Horizontal lines within the box plots represent the median values (*** p<0.001).

## Discussion

As a mannitol stress assay did not result in our hands in statistically significant phenotypic differences (Neves *et al*., 2021, 2022; Waese *et al*., 2017) we used *Pseudomonas* infection assays to address possible biotic stress functional context for EXO70E2 exocyst subunit. Using flooding assays in Arabidopsis WT vs. *exo70E2* mutant we identified significant enhanced sensitivity of the mutant to both WT and T3SS mutant of *Pseudomonas syringae* attack. This is in agreement with recently published observations showing enhanced *exo70E2 LOF* mutant sensitivity to hemibiotrophic fungus *Colletotrichum higginsianum* (Koch *et al*., 2026) and EXO70E2 overexpressing plants’ enhanced sensitivity to fungal necrotroph *Botrytis cinerea* (Wu and Huang, 2026). Correlating with the enhanced sensitivity we have observed lower callose accumulation in response to the *Pseudomonas* attack as did Reddit et al. (Redditt *et al*., 2019) in response to flagellin treatment in the *exo70E2* mutant. These observations clearly point to the plant defence processes - both to bacteria as well as fungi - as the EXO70E2 functional integration framework. Moreover - our and data published before show that EXO70E2 interacts with several core exocyst subunits (Fig. S2; Ding et al., 2013) so that it is reasonable to assume that EXO70E2 functions as a subunit of a specific form of the exocyst complex (Zárský *et al*., 2013).

EXO70E2 was, despite original interpretations (see the Introduction), linked to the selective autophagy via NBR1 effector interaction (Ji *et al*., 2020; Yan *et al*., 2024). However, autophagy flux in the *exo70E2* LOF mutant - basal and after the pathogen infection - was not studied yet. In the *nbr1* mutant background accumulation of EXO70E2-GFP into the vacuole was diminished by about 50% (Ji *et al*., 2020), correlating well with the established function of NBR1 as an autophagy receptor in EXO70E2 loading to the autophagosomes (Ji *et al*., 2020; Yan *et al*., 2024). It is interesting that despite different autophagy boosting treatments (BTH in Ji *et al.,* 2020 and *Pseudomonas* infection here in our experiments) we observed that the effect of loss of EXO70E2 function downregulates NBR1 delivery to the vacuole to a very comparable proportion (Fig. 3), supporting the notion that based on a direct (and probably stoichiometric - see the model in Yan *et al.,* 2024) interaction EXO70E2 is an important component of NBR1-dependent autophagic flux. However, since *P. syringae* infection has been reported to induce NBR1 expression (Üstün et *al.,* 2018), altered NBR1 expression in the *exo70E2* mutant cannot currently be excluded. We have seen that *Pseudomonas* induces protein accumulation of both NBR1 and EXO70E2 (Figure S1). Nevertheless, given the established NBR1 - EXO70E2 direct interaction, altered NBR1 turnover represents a more direct mechanistic explanation. As an alternative approach, experiment with the block of part of the autophagy cargo degradation using E-64d protease inhibitor, indicates - both in microscopy compartment accumulation as well as in Western blot analysis - that also basal autophagy pathway under normal control conditions is downregulated in the *exo70E2* LOF mutant, similarly e.g. to the situation in a loss of core autophagy pathway regulators ATG2 or ATG5 (but not in *atg9* mutant; (Inoue *et al*., 2006). Also, Goto-Yamada et al. (2019) observed different mutants defective in ATG2, ATG5, and ATG7 treated with E-64d, and showed varying degrees of reduction in the size of E-64d compartments. Both tests of autophagy cargo accumulation after the ConA or E-64d treatments blocking degradation show distinct downregulation of autophagy flux in *exo70E2* mutant which could be only partially explained as the loss of NBR1 receptor’s EXO70E2 protein cargo. Distinctly decreased FM4-64 visualized compartments accumulation after the E-64d treatment in *exo70E2* mutant vs WT indicate that EXO70E2 possibly contributes more to a general autophagy process than just being a cargo for selective autophagy. For instance, it might be that NBR1-EXO70E2 substantially affects liquid-liquid phase transition dynamics of droplet autophagosome precursors (Yan *et al.,* 2024). While the direct role in autophagy-related processes remains a parsimonious interpretation of our data, at the moment we cannot exclude other roles of EXO70E2 that explain observed differences in autophagic flux. EXO70E2 may influence NBR1-dependent vacuolar trafficking indirectly, for example through effects on endomembrane dynamics, vacuolar function, or the delivery of components required for efficient degradation. In addition, although NBR1 vacuolar delivery is generally ATG-dependent and associated with canonical macroautophagy, recent evidence for NBR1 involvement in microautophagy-like processes suggests that EXO70E2 may also affect non-canonical routes of vacuolar cargo delivery (Lee *et al*., 2023). Further mechanistic studies will be required to distinguish between these possibilities. Finally, we have to note that while we could see GFP-EXO70E2 fluorescence in the root, the *Pseudomonas* growth, callose deposition, and NBR1 detection upon flooding were all assayed in the shoot. According to publicly available databases (ePlant - University of Toronto, Waese *et al*., 2017), EXO70E2 is expressed in the shoot (esp. stomata) as well, but our natural EXO70E2 promoter::GFP-EXO70E2-complementing construct does not show visible signal. Nevertheless, our data collectively point at involvement of EXO70E2 in regulation of selective autophagic flux. Results presented here support the notion that Arabidopsis exocyst subunits are not only cargo of (selective) autophagy but may also engage in regulation of autophagic flux. Further research is needed to determine the exact step at which EXO70E2 regulates autophagic flux.

## Supplementary data

Fig. S1. Western blot analysis of e2 Comp seedlings under Pst Dc3000 infection

Fig. S2. Yeast two-hybrid (Y2H) analysis of interaction between EXO70E2 and other exocyst subunits.

Fig. S3. Root length measurement of *exo70* mutants subjected to mannitol stress.

## Acknowledgements

We thank Dr. Tamara Pečenková for help with designing the *Pseudomonas* flux assay. We would like to thank Dr. Jitka Ortmannová for advice and discussions on autophagy related experiments. Microscopy works were performed at the Imaging Facility of the IEB AS CR, supported by MEYS CR LM2023050 ‘Czech-BioImaging’ and IEB AS CR.

## Author contributions

A.B.Y. performed most of the experiments, A.P. and A.B.Y. did *Pseudomonas* assays, A.B. prepared and verified the *exo70E2* CRISPR line, P.S. and G.A.C. contributed to planning of the project, interpretation of data and writing, and V. Ž. planned the project was responsible for project coordination, experimental design and interpretation of data.

## Data availability

Raw data for experiments where not shown is available upon request.

## References

Acheampong AK, Shanks C, Cheng C-Y, Schaller GE, Dagdas Y, Kieber JJ. 2020. EXO70D isoforms mediate selective autophagic degradation of type-A ARR proteins to regulate cytokinin sensitivity. Proceedings of the National Academy of Sciences of the United States of America 117, 27034–27043.

Allen H, Davis B, Patel J, Gu Y. 2024. Dot Scanner: open-source software for quantitative live-cell imaging in planta. The Plant Journal: For Cell and Molecular Biology 118, 1689–1698.

Bodemann BO, Orvedahl A, Cheng T, et al. 2011. RalB and the exocyst mediate the cellular starvation response by direct activation of autophagosome assembly. Cell 144, 253–267.

Brillada C, Teh O-K, Ditengou FA, et al. 2021. Exocyst subunit Exo70B2 is linked to immune signaling and autophagy. The Plant Cell 33, 404–419.

Camacho L. 2019. Characterization of the interaction between ISTL1 and Exo70E2 in Arabidopsis. FASEB Journal: Official Publication of the Federation of American Societies for Experimental Biology 33.

Clough SJ, Bent AF. 1998. Floral dip: a simplified method for Agrobacterium-mediated transformation of Arabidopsis thaliana. The Plant Journal: For Cell and Molecular Biology 16, 735–743.

Copeland C, Logemann E, Malisic M, Amrhein A, Valisi A, Schulze-Lefert P. 2025. A multigenic quantitative trait locus underlies natural variation in *Arabidopsis thaliana* root system architecture and transcriptional responses to microbiota-derived *Pseudomonas*. bioRxiv.

Cvrčková F, Grunt M, Bezvoda R, Hála M, Kulich I, Rawat A, Zárský V. 2012. Evolution of the land plant exocyst complexes. Frontiers in Plant Science 3, 159.

Cvrčková F, Zárský V. 2013. Old AIMs of the exocyst: evidence for an ancestral association of exocyst subunits with autophagy-associated Atg8 proteins. Plant Signaling & Behavior 8, e27099.

Dikic I. 2017. Proteasomal and autophagic degradation systems. Annual Review of Biochemistry 86, 193–224.

Ding Y, Wang J, Chun Lai JH, et al. 2014. Exo70E2 is essential for exocyst subunit recruitment and EXPO formation in both plants and animals. Molecular Biology of the Cell 25, 412–426.

Drs M, Müller K, Voxeur A, Ortmannová J, Serrano N, Vernhettes S, Škrabálková E, Potocký M, Žárský V, Pečenková T. 2025. Exocyst subunits EXO70B1 and B2 contribute to stomatal dynamics and cell wall modifications. Frontiers in Plant Science 16, 1694769.

Du Y, Mpina MH, Birch PRJ, Bouwmeester K, Govers F. 2015. Phytophthora infestans RXLR effector AVR1 interacts with exocyst component Sec5 to manipulate plant immunity. Plant Physiology 169, 1975–1990.

Du Y, Overdijk EJR, Berg JA, Govers F, Bouwmeester K. 2018. Solanaceous exocyst subunits are involved in immunity to diverse plant pathogens. Journal of Experimental Botany 69, 655–666.

Engler C, Kandzia R, Marillonnet S. 2008. A one pot, one step, precision cloning method with high throughput capability. PloS One 3, e3647.

Farré J-C, Subramani S. 2016. Mechanistic insights into selective autophagy pathways: lessons from yeast. Nature Reviews. Molecular Cell Biology 17, 537–552.

Goto-Yamada S, Oikawa K, Bizan J, et al. 2019. Sucrose starvation induces microautophagy in plant root cells. Frontiers in Plant Science 10, 1604.

Hafrén A, Üstün S, Hochmuth A, Svenning S, Johansen T, Hofius D. 2018. Turnip mosaic virus counteracts selective autophagy of the viral silencing suppressor HCpro. Plant Physiology 176, 649–662.

Hála M, Cole R, Synek L, et al. 2008. An exocyst complex functions in plant cell growth in Arabidopsis and tobacco. The Plant Cell 20, 1330–1345.

Holden S, Bergum M, Green P, et al. 2022. A lineage-specific Exo70 is required for receptor kinase-mediated immunity in barley. Science Advances 8, eabn7258.

Inoue Y, Suzuki T, Hattori M, Yoshimoto K, Ohsumi Y, Moriyasu Y. 2006. AtATG genes, homologs of yeast autophagy genes, are involved in constitutive autophagy in Arabidopsis root tip cells. Plant & Cell Physiology 47, 1641–1652.

Ishiga Y, Ishiga T, Uppalapati SR, Mysore KS. 2011. Arabidopsis seedling flood-inoculation technique: a rapid and reliable assay for studying plant-bacterial interactions. Plant Methods 7, 32.

Ji C, Zhou J, Guo R, Lin Y, Kung C-H, Hu S, Ng WY, Zhuang X, Jiang L. 2020. AtNBR1 is a selective autophagic receptor for AtExo70E2 in Arabidopsis. Plant Physiology 184, 777–791.

Koch BL, Gardner D, Smith H, et al. 2026. Molecular insights into the production of extracellular vesicles by plants. Plant Physiology 200.

Koncz C, Schell J. 1986. The promoter of TL-DNA gene 5 controls the tissue-specific expression of chimaeric genes carried by a novel type of Agrobacterium binary vector. Molecular & General Genetics 204, 383–396.

Kubalová M, Griffiths J, Müller K, Levenets L, Tylová E, Tarkowská D, Jones AM, Fendrych M. 2025. Gibberellin-deactivating GA2OX enzymes act as a hub for auxin-gibberellin cross talk in Arabidopsis thaliana root growth regulation. Proceedings of the National Academy of Sciences of the United States of America 122, e2425574122.

Kulich I, Pečenková T, Sekereš J, Smetana O, Fendrych M, Foissner I, Höftberger M, Zárský V. 2013. Arabidopsis exocyst subcomplex containing subunit EXO70B1 is involved in autophagy-related transport to the vacuole. Traffic (Copenhagen, Denmark) 14, 1155–1165.

Kulich I, Vojtíková Z, Glanc M, Ortmannová J, Rasmann S, Žárský V. 2015. Cell wall maturation of Arabidopsis trichomes is dependent on exocyst subunit EXO70H4 and involves callose deposition. Plant Physiology 168, 120–131.

Kulich I, Vojtíková Z, Sabol P, Ortmannová J, Neděla V, Tihlaříková E, Žárský V. 2018. Exocyst subunit EXO70H4 has a specific role in callose synthase secretion and silica accumulation. Plant Physiology 176, 2040–2051.

Lee HN, Chacko JV, Gonzalez Solís A, Chen K-E, Barros JAS, Signorelli S, Millar AH, Vierstra RD, Eliceiri KW, Otegui MS. 2023. The autophagy receptor NBR1 directs the clearance of photodamaged chloroplasts. eLife 12, e86030.

Lei Y, Lu L, Liu H-Y, Li S, Xing F, Chen L-L. 2014. CRISPR-P: a web tool for synthetic single-guide RNA design of CRISPR-system in plants. Molecular Plant 7, 1494–1496.

Lin Y, Ding Y, Wang J, et al. 2015. Exocyst-positive organelles and autophagosomes are distinct organelles in plants. Plant Physiology 169, 1917–1932.

Lin Y, Zeng Y, Zhu Y, Shen J, Ye H, Jiang L. 2021. Plant Rho GTPase signaling promotes autophagy. Molecular Plant 14, 905–920.

Marshall RS, Vierstra RD. 2018. Autophagy: The master of bulk and selective recycling. Annual Review of Plant Biology 69, 173–208.

Michalopoulou VA, Mermigka G, Kotsaridis K, Mentzelopoulou A, Celie PHN, Moschou PN, Jones JDG, Sarris PF. 2022. The host exocyst complex is targeted by a conserved bacterial type-III effector that promotes virulence. The Plant Cell 34, 3400–3424.

Mostowy S. 2013. Autophagy and bacterial clearance: a not so clear picture. Cellular Microbiology 15, 395–402.

Neves J, Monteiro J, Sousa B, Soares C, Pereira S, Fidalgo F, Pissarra J, Pereira C. 2022. Relevance of the exocyst in Arabidopsis exo70e2 mutant for cellular homeostasis under stress. International Journal of Molecular Sciences 24, 424.

Neves J, Sampaio M, Séneca A, Pereira S, Pissarra J, Pereira C. 2021. Abiotic stress triggers the expression of genes involved in Protein Storage Vacuole and exocyst-mediated routes. International Journal of Molecular Sciences 22, 10644.

Ortmannová J, Sekereš J, Kulich I, Šantrůček J, Dobrev P, Žárský V, Pečenková T. 2022. Arabidopsis EXO70B2 exocyst subunit contributes to papillae and encasement formation in antifungal defence. Journal of Experimental Botany 73, 742–755.

Pecenková T, Hála M, Kulich I, Kocourková D, Drdová E, Fendrych M, Toupalová H, Zársky V. 2011. The role for the exocyst complex subunits Exo70B2 and Exo70H1 in the plant-pathogen interaction. Journal of Experimental Botany 62, 2107–2116.

Pecenková T, Janda M, Ortmannová J, Hajná V, Stehlíková Z, Žárský V. 2017a. Early Arabidopsis root hair growth stimulation by pathogenic strains of Pseudomonas syringae. Annals of Botany 120, 437–446.

Pecenková T, Markovic V, Sabol P, Kulich I, Žárský V. 2017b. Exocyst and autophagy-related membrane trafficking in plants. Journal of Experimental Botany 69, 47–57.

Pečenková T, Potocká A, Potocký M, et al. 2020. Redundant and diversified roles among selected Arabidopsis thaliana EXO70 paralogs during biotic stress responses. Frontiers in Plant Science 11, 960.

Redditt TJ, Chung E-H, Zand Karimi H, et al. 2019. AvrRpm1 functions as an ADP-ribosyl transferase to modify NOI-domain containing proteins, including Arabidopsis and soybean RPM1-interacting protein 4. The Plant Cell 31, 2664–2681.

Serre NBC, Kralík D, Yun P, Slouka Z, Shabala S, Fendrych M. 2021. AFB1 controls rapid auxin signalling through membrane depolarization in Arabidopsis thaliana root. Nature Plants 7, 1229–1238.

Stegmann M, Anderson RG, Ichimura K, Pecenkova T, Reuter P, Žársky V, McDowell JM, Shirasu K, Trujillo M. 2012. The ubiquitin ligase PUB22 targets a subunit of the exocyst complex required for PAMP-triggered responses in Arabidopsis. The Plant Cell 24, 4703–4716.

Svenning S, Lamark T, Krause K, Johansen T. 2011. Plant NBR1 is a selective autophagy substrate and a functional hybrid of the mammalian autophagic adapters NBR1 and p62/SQSTM1. Autophagy 7, 993–1010.

Synek L, Pleskot R, Sekereš J, et al. 2021. Plasma membrane phospholipid signature recruits the plant exocyst complex via the EXO70A1 subunit. Proceedings of the National Academy of Sciences of the United States of America 118, e2105287118.

Synek L, Schlager N, Eliás M, Quentin M, Hauser M-T, Zárský V. 2006. AtEXO70A1, a member of a family of putative exocyst subunits specifically expanded in land plants, is important for polar growth and plant development. The Plant Journal: For Cell and Molecular Biology 48, 54–72.

Teh O-K, Lee C-W, Ditengou FA, et al. 2018. Phosphorylation of the exocyst subunit Exo70B2 contributes to the regulation of its function. bioRxiv.

TerBush DR, Maurice T, Roth D, Novick P. 1996. The Exocyst is a multiprotein complex required for exocytosis in Saccharomyces cerevisiae. The EMBO Journal 15, 6483–6494.

Tzfadia O, Galili G. 2013. The Arabidopsis exocyst subcomplex subunits involved in a golgi-independent transport into the vacuole possess consensus autophagy-associated atg8 interacting motifs. Plant Signaling & Behavior 8, doi: 10.4161/psb.26732.

Üstün S, Hafrén A, Liu Q, Marshall RS, Minina EA, Bozhkov PV, Vierstra RD, Hofius D. 2018. Bacteria exploit autophagy for proteasome degradation and enhanced virulence in plants. The Plant Cell 30, 668–685.

Waese J, Fan J, Pasha A, et al. 2017. EPlant: Visualizing and exploring multiple levels of data for hypothesis generation in plant biology. The Plant Cell 29, 1806–1821.

Wang J, Ding Y, Wang J, Hillmer S, Miao Y, Lo SW, Wang X, Robinson DG, Jiang L. 2010. EXPO, an exocyst-positive organelle distinct from multivesicular endosomes and autophagosomes, mediates cytosol to cell wall exocytosis in Arabidopsis and tobacco cells. The Plant Cell 22, 4009–4030.

von Wangenheim D, Hauschild R, Fendrych M, Barone V, Benková E, Friml J. 2017. Live tracking of moving samples in confocal microscopy for vertically grown roots. eLife 6, e26792.

Wang W, Liu N, Gao C, Cai H, Romeis T, Tang D. 2020. The Arabidopsis exocyst subunits EXO70B1 and EXO70B2 regulate FLS2 homeostasis at the plasma membrane. The New Phytologist 227, 529–544.

Wickham H, Averick M, Bryan J, et al. 2019. Welcome to the tidyverse. Journal of Open Source Software 4, 1686.

Wu X, Huang J. 2026. Overexpression of EXO70E2 in Arabidopsis thaliana Disrupts Normal Development and Enhances Susceptibility to the Necrotrophic Fungus Botrytis cinerea. Genes 17, 347.

Xing H-L, Dong L, Wang Z-P, Zhang H-Y, Han C-Y, Liu B, Wang X-C, Chen Q-J. 2014. A CRISPR/Cas9 toolkit for multiplex genome editing in plants. BMC Plant Biology 14, 327.

Yan H, Qi A, Lu Z, et al. 2024. Dual roles of AtNBR1 in regulating selective autophagy via liquid-liquid phase separation and recognition of non-ubiquitinated substrates in Arabidopsis. Autophagy 20, 2804–2815.

Zaffagnini G, Savova A, Danieli A, et al. 2018. Phasing out the bad-How SQSTM1/p62 sequesters ubiquitinated proteins for degradation by autophagy. Autophagy 14, 1280–1282.

Zárský V, Kulich I, Fendrych M, Pečenková T. 2013. Exocyst complexes multiple functions in plant cells secretory pathways. Current Opinion in Plant Biology 16, 726–733.

Žárský V, Sekereš J, Kubátová Z, Pečenková T, Cvrčková F. 2020. Three subfamilies of exocyst EXO70 family subunits in land plants: early divergence and ongoing functional specialization. Journal of Experimental Botany 71, 49–62.

Zhou F, Wu Z, Zhao M, Segev N, Liang Y. 2019. Autophagosome closure by ESCRT: Vps21/RAB5-regulated ESCRT recruitment via an Atg17-Snf7 interaction. Autophagy 15, 1653–1654.

Zhou J, Zhang Y, Qi J, Chi Y, Fan B, Yu J-Q, Chen Z. 2014. E3 ubiquitin ligase CHIP and NBR1-mediated selective autophagy protect additively against proteotoxicity in plant stress responses. PLoS Genetics 10, e1004116.

