## Supplementary figures to E2 for "Arabidopsis exocyst complex subunit EXO70E2 in defence against *Pseudomonas syringae* in conjunction with autophagy"

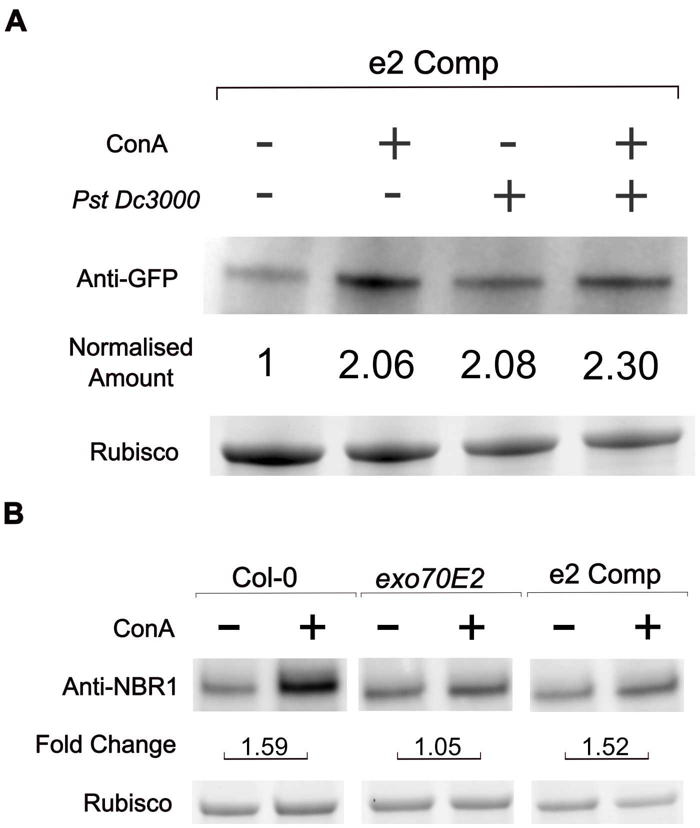


**Fig. S1.** Western blot analysis of e2 Comp seedlings under *Pst Dc3000* infection. (A)Western blot analysis of GFP tagged EXO70E2 protein in 7-day-old e2 Comp Arabidopsis seedlings. Seedlings were subjected to bathing inoculation with *Pst Dc3000*, in the presence (+) or absence (–) of ConA at 10 hpi for additional 14h. Total protein extracts were immunoblotted with anti-GFP antibodies overnight. Intesity of anti-GFP bands were first normalised according to the load (Rubisco) and mock condition ( *Pst Dc3000* (-) and ConA(-)) was normalised to 1 and the intensity of other conditions were calculated according to mock condition. (B)Western blot analysis of NBR1 protein accumulation and autophagic flux in 11-day-old wild-type (Col-0), *exo70E2* mutant and e2 Comp Arabidopsis seedlings. Seedlings were subjected to flooding inoculation with *Pst Dc3000*, in the presence (+) or absence (–) of ConA at 10 hpi for additional 14h. Total protein extracts were immunoblotted with anti-NBR1 antibodies overnight. Rubisco (Bio-Rad stain-free gel activation) serves as the loading control. Fold-change values indicate the ratio of NBR1 band intensity between ConA(+) and (-) conditions for each respective genotype after the *Pst* infection.

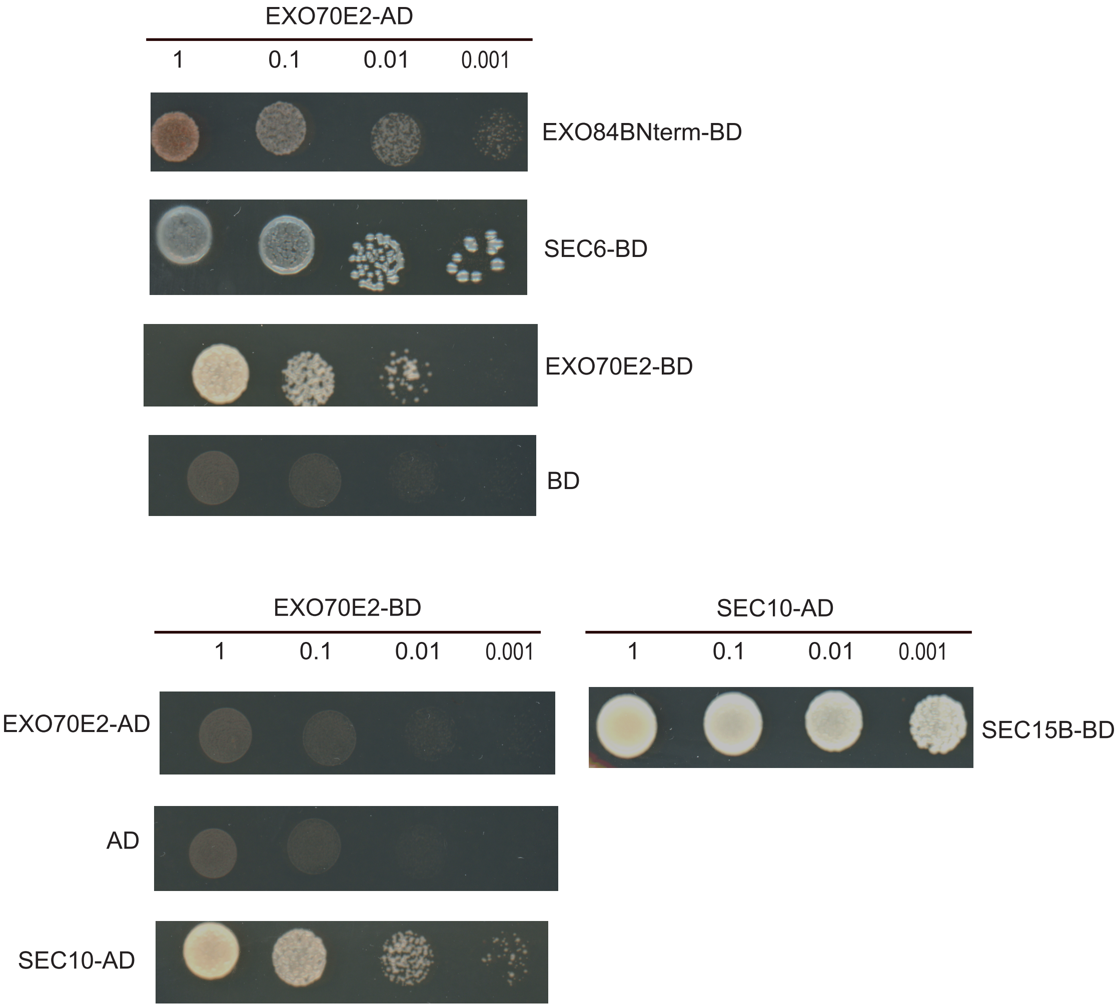


**Fig. S2**. Yeast two-hybrid (Y2H) analysis of interaction between EXO70E2 and other exocyst subunits. *S. cerevisiae* cultures expressing recombinant EXO70E2 and other exocyst subunit constructs were subjected to a 10-fold serial dilution assay (1, 0.1, 0.01, and 0.001, from left to right) to assess interaction strength. SEC10-AD and SEC15B-BD interaction was used as a positive control. Growth assays were performed on synthetic defined (SD) medium lacking Adenine, Histidine, Leucine, and Tryptophan(–Ade/–His–Leu/–Trp) to select for reporter gene activation.

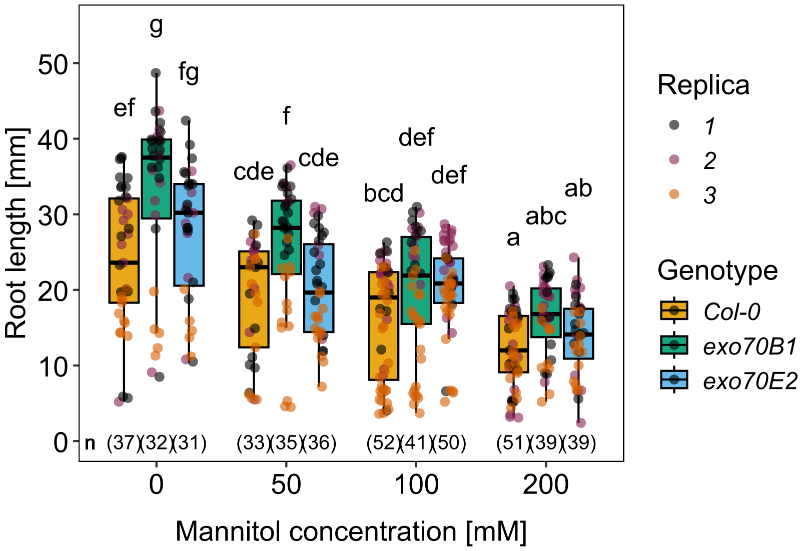


**Fig. S3.** Root length measurement of *exo70e2* mutants after subjected to osmotic stress. Arabidopsis seedlings were grown for 9 days on half MS (1/2 MS) 1 % agar plates or 1/2 MS 1 % agar plates supplemented with indicated concentration of mannitol. Shown are median values of primary root lengths. Data were analyzed using a linear model with Genotype, Treatment, and their interaction as fixed effects, and Replica as an additional fixed effect to account for differences among experimental replicates. The significance of model terms was assessed by analysis of variance (ANOVA). Estimated marginal means (EMMs) were calculated for each genotype × treatment combination, and pairwise comparisons were performed using Tukey's honestly significant difference (HSD) adjustment for multiple testing. The interaction analysis showed no significant effect of *exo70e2* genotype..
